# Transcription factors associated with regulation of transcriptome in human thigh and calf muscles at baseline and after six days of disuse

**DOI:** 10.1101/2024.12.10.627667

**Authors:** Anna A. Borzykh, Pavel A. Makhnovskii, Ivan I. Ponomarev, Tatiana F. Vepkhvadze, Egor M. Lednev, Ilya V. Rukavishnikov, Oleg I. Orlov, Elena S. Tomilovskaya, Daniil V. Popov

**Affiliations:** Institute of Biomedical Problems of the Russian Academy of Sciences, Moscow, Russia

**Keywords:** dry immersion, skeletal muscle, RNA-seq, position weight matrix, transcription factor binding site, transcription factor

## Abstract

Disuse has a negative impact on the postural muscles of the trunk and legs. Different leg muscles demonstrate a differentiated and conservative response to disuse, in terms of a decrease in muscle mass, strength, aerobic performance, and changes in gene expression. We aimed to identify transcription factors regulating gene expression at baseline and after disuse in human *m. soleus* – a “slow” muscle with a strong postural function, and “mixed” *m. vastus lateralis*. Biopsies were taken from these muscles prior to and after 6 days of strict disuse (dry immersion). The enriched transcription factor binding sites (and corresponding factors) in the individual promoter regions of co-expressed genes were examined using the positional weight matrix approach. The baseline transcriptomic profiles and the disuse-induced changes (RNA-seq) differ significantly between muscles. In particular, the specific and significant response to disuse in *m. soleus* was found to be strongly related to the suppression of genes regulating the mitochondrial energy metabolism, the activation of the inflammatory response and the ubiquitin-proteasome system. This response is associated with the proinflammatory transcription factors such as families IRF, STAT, and other. The validity of approximately two-thirds of the predicted transcription factors was indirectly confirmed by the analysis of their function described in the literature. These identified transcription factors appear to be promising candidates for future targeted studies that mechanistically investigate gene expression regulation in various muscles at baseline, following disuse or inactivity.

**Highlights:**

- Disuse has a different negative impact on the different human postural leg muscles.
- The transcriptome regulation in *m. soleus* and *m. vastus lateralis* differs markedly.
- The gene response to disuse in *m. soleus* is greater than in *m. vastus lateralis*.
- This partially related to activation of inflammation-induced transcription factors.

## Introduction

A significant decrease in the mass and functional capabilities of skeletal muscles: strength [1], aerobic performance (endurance) [2], and insulin sensitivity [3], is occurred in the first weeks of disuse (limb immobilization, bedrest, dry immersion, etc.). This problem is relevant for people in bed rest (e.g., with injuries of the musculoskeletal system and other pathological conditions), as well as during prolonged stay in confined spaces (e.g., spaceflight, pandemic isolation). Therefore, the investigation of the mechanisms underlying the reduction in muscle mass and functional capacity during disuse/inactivity, and the development of approaches (particularly pharmacological) to prevent these changes, is both a fundamental and important applied challenge.

Disuse primarily causes changes in the postural muscles of the trunk and legs, in particular, in *m. soleus* – a “slow” calf muscle with a pronounced postural function. Differentiated response of different leg muscles to disuse/spaceflight is observed in different species: in humans [4], primates [5, 6], rats [7] and mice [8-10], which indicates the conservative mechanism its regulation.

Apparently, the differentiated response to disuse in different muscles is associated with differences in their daily activity, neural control [11], and the muscle fiber type composition. Thus, human *m. soleus*, unlike other leg muscles, contains more than 80% slow muscle fibers [12], which correlates with pronounced differences in the transcriptomic and proteomic profiles between *m. soleus* and other leg muscles or between slow- and fast-twitch muscle fibers [13-16]. In humans, the disuse-induced decrease in the mass of the “mixed“ (about 45% slow-twitch fibers) *m. vastus lateralis* is mainly regulated by a decrease in the translation rate [17, 18], while the role of the ubiquitin-proteasome system, which plays a key role in proteolysis in skeletal muscle, is yet unclear [19]. However, in rodents, disuse causes activation of proteolysis specific to “slow” *m. soleus* [20]. In addition, disuse/spaceflight-induced transcriptome changes in *m. soleus* in humans after 7-day dry immersion [16] and 2-month bedrest [21], as well as in rodents after 1-week hindlimb suspension [7] and 1-month spaceflight [8, 9]) are several times more pronounced than in other leg muscles. This suggests that regulation at the transcriptional level may also play a role in disuse-induced decrease in muscle mass and functional capacity, in particular in *m. soleus*. However, the regulation of this differentiated gene (transcriptomic) response to disuse in different muscles remains unexplored.

It should be noted that the search for transcription factors (TFs) regulating the expression of hundreds (sometimes even thousands) of genes is a very complex task. To solve this problem, the system analyzed in this work was simplified, namely, TFs regulating expression was searched in clusters of co-expressed (co-regulated) genes; clusters were identified using differences in *i*) transcriptome profiles between *m. soleus* and *m. vastus lateralis* at baseline and *ii*) transcriptome response to a strict disuse – 6 days of dry immersion. To search for TFs, the classical approach – the positional weight matrix (PWM) approach, was used. Unlike previous studies investigating transcriptome regulation in human skeletal muscle, enrichment of TF binding sites was examined in individual promoter regions defined for human skeletal muscle by the position of transcription start site (CAGE-seq) and surrounding open chromatin (DNase-seq and ATAC-seq) [22].

## Methods

### Ethical approval

The study was approved by the Committee on Biomedical Ethics of the Institute of Biomedical Problems of the RAS (protocol No. 594 of September 6, 2021) and complied with all the principles set forth in the Declaration of Helsinki. All volunteers gave written informed consent to participate in the study and to undergo skeletal muscle biopsy.

### Study design

Design of the study was described previously [16]. Briefly, ten males participated in a 7-day dry immersion (DI, ages 25–38 years; median height 1.78 m [interquartile range 1.71–1.79 m]; body mass 70 kg [64–74 kg]; and body mass index 24 kg/m^2^ [21–24 kg/m^2^]). Muscle samples were taken using a Bergström needle with aspiration under local anesthesia (2 ml of 2% lidocaine) from the medial part of *m. vastus lateralis* (VL) and *m. soleus* (S) muscles of the left leg 14 days before the start of DI (prior to any familiarization protocols and physiological tests, which did not included in the paper) and 6 days after the start of 7-day DI (at 10:00, 3 hours after a standardized light breakfast: 5.2 g protein, 2.7 g fat, 55 g carbohydrates, 1253 kJ); this approach allowed us to study the “pure” gene response to 6 days of disuse. The samples were cleared of visible fragments of connective and adipose tissue, frozen in liquid nitrogen, and stored at -80 ºC.

### RNA sequencing and data processing

A workflow was described previously [16]: a piece of frozen tissue (∼15 mg) was homogenized at +4 °C in 500 μl of a lysis buffer (ExtractRNA, Evrogen, Russia) using a plastic pestle and a drill. Total RNA was isolated using spin columns (RNeasy Mini Kit, Qiagen, Germany). RNA concentration was assessed using a Qubit 4 fluorimeter (ThermoScientific, USA); RNA integrity – using capillary gel electrophoresis (TapeStation, Agilent, USA). All samples had RNA integrity (RIN) >7.8. Strand-specific RNA libraries were prepared using the NEBNext Ultra II RNA kit (New England Biolabs, USA) and single-end sequencing was done by a NextSeq 550 analyzer (Illumina, USA) with a read length of 75 bp (∼60 million reads/sample), as described previously [23].

Data processing was described previously [16]: sequencing quality was assessed using the FastQC tool (v.0.11.5, RRID:SCR_014583), and low-quality reads were removed using Trimmomatic (v.0.36, RRID:SCR_011848). High-quality reads were mapped to the human reference genome GRCh38.p13 primary assembly. The number of unique reads mapped to exons of each gene was determined using the Rsubread package (R environment) and Ensembl annotation (GRCh38.101). The DESeq2 method (paired sample analysis with Benjamini-Hochberg correction) was used to analyze differential gene expression. Differentially expressed genes were defined as protein-coding genes (as well as polymorphic pseudogenes and translated pseudogenes) with an expression level of TPM >1 (Transcripts Per Million, kallisto tool v0.46.2), *p*_*adj*_<0.01 and |fold change|≥1.25.

### Functional enrichment analysis

Enrichment analysis (biological processes, cellular components and pathways) was performed against all expressed protein-coding genes (TPM >1, ∼10,000 mRNA) by the DAVID tool v2023q4 (RRID:SCR_001881) using UNIPROT KW BP/CC, GENE ONTOLOGY BP/CC DIRECT and KEGG PATHWAY databases, Fisher’s exact test with Benjamini correction, and *p*_*adj*_<0.05.

### Clusters of co-expressed genes

Expression values (normalized read counts) for each gene (4 experimental points [2 time points and 2 muscles] × 10 subjects) were ranked. The co-expressed genes were identified by an unsupervised analysis (the Chinese restaurant process [24]) using the geneXplain platform (function “CRC clustering,” cluster process number□=□100, cycles per clustering process□=□100, without considering inverted profiles as similar) (http://wiki.biouml.org/index.php/CRC_Analysis).

Then group of clusters with similar patterns of gene regulation in *i*) VL and S at baseline (e.g., ‘*Up-regulated in S*’) and *ii*) following disuse (e.g., ‘*Down-regulated in S only*’) were identified using Euclidean distance (the Ward’s method).

### Putative transcription factors (PWM method)

Using the PWM method, the transcription factor binding sites (TFBSs) (and corresponding TFs) were predicted in the individual promoter regions for each cluster. The position of the individual promoter regions in human skeletal muscle was evaluated previously using data of the transcription start site [CAGE-seq] and the surrounding open chromatin [DNase-seq and ATAC-seq]) [22]. Enrichment of predicted TFBSs were performed by the geneXplain platform (the “Search for enriched TFBSs (tracks)” function http://wiki.biouml.org/index.php/Search_for_enriched_TFBSs_(tracks)_(analysis)) using the PWM database TRANSFAC v2022.2 [25, 26]. Namely, the maximum enrichment (FE_adj_, statistically corrected odds ratios with a confidence interval of 99%) was determined for each PWM relative to that in 5000 random individual promoters showing no differential expression in any of experimental time points (DESeq2 method, *p*_*adj*_□>□0.5). Adjusted fold enrichment (FE_adj_)□>□1.5 for transcription factor binding site (the binomial test) and FDR□<□0.05 were set as significance thresholds. If a TF has several PWM, the most enriched PWM was used.

### Validation of putative transcription factors (Medline search)

To validate TFs identified in our study associations of these TFs with enriched biological term/function (UNIPROT KW BP/CC) in muscle or other tissues were exanimated in each cluster by a Medline search. For the Medline search, terms: *TF symbol* OR *TF family name* (from the HOCOMOCO v12 database) AND *enriched term for co-regulated genes* AND *muscle* (from Figure 3 and 4) were used. Articles showing a link between predicted TF and enriched biological function in muscle or other tissues were selected.

## Results and Discussion

### Difference in transcriptomic profiles in *m. vastus lateralis* and *m. soleus* at baseline and in their response to 6 days of disuse

As described elsewhere [16], after the removal of low-expressed genes about 10,000 mRNAs were detected (Supporting information Table S1). It was found that 1341 mRNAs up-regulated in VL compared to S at baseline enriched terms related to translation and transcription, glycolytic enzymes, and endoplasmic reticulum/calcium signaling. Meanwhile, 1152 mRNAs down-regulated in VL (or up-regulated in S) enriched terms related to aerobic and fatty acid metabolism enzymes, contractile proteins, cell adhesion, and ion transporters (Fig. 1A and B, Supporting information Table S1 and S2).

**Fig. 1.**
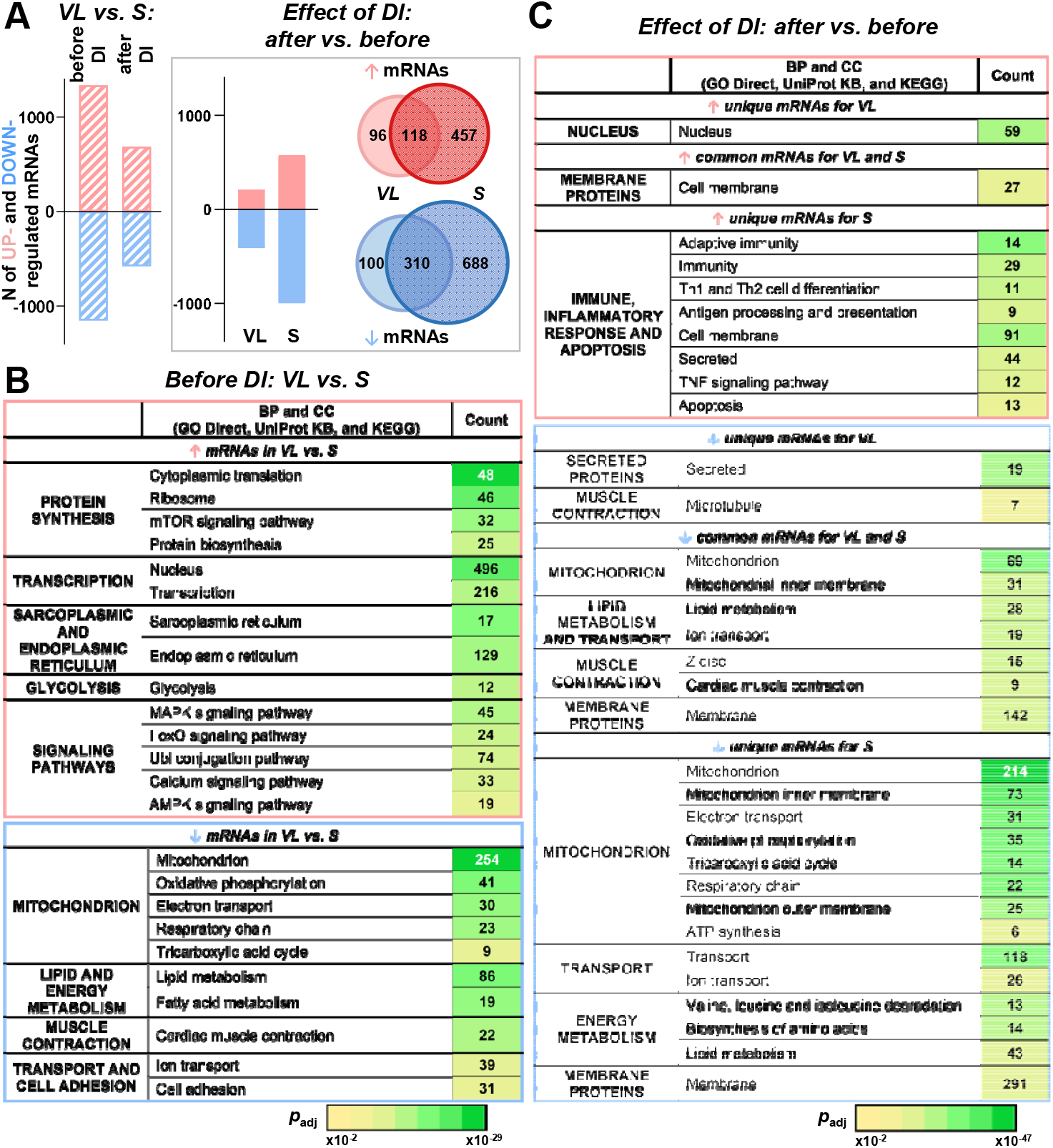
*M. vastus lateralis* (VL) and *m. soleus* (S) demonstrated pronounced differences in transcriptome at baseline (prior to dry immersion) and changes in transcriptome induced by 6 days of dry immersion. A – Number up- and down-regulated mRNAs in VL *vs*. S at baseline and following dry immersion, as well as in each muscle following dry immersion (see a list of all detected protein-coding genes in Supporting information Table S1; *n* = 10 subjects). B and C – Main result of functional enrichment analysis (all results are presented in Supporting information Table S2) for VL *vs*. S at baseline (B) and for unique and common genes those response to dry immersion in VL and S (C). The number of genes in each category is indicated, the heat map shows *p*_*adj*_.

Disuse-induced changes in gene expression in S were 2.5 times (∼1500 mRNAs) more pronounced than in VL (∼600 mRNAs). Interestingly, only partial overlap in gene response was found between muscles (Fig. 1A) that indicates muscle-specific regulation of gene response to disuse. Functional enrichment analysis showed that following disuse, both muscles demonstrated decreased expression of genes encoding numerous membrane proteins, mitochondrial energy metabolism enzymes, and several sarcomeric proteins. In S, a decrease in the expression of a significantly larger number of genes encoding mitochondrial and membrane (especially transport) proteins was found, as well as an increase in the expression of inflammatory genes (Fig. 1C).

Additionally, the more pronounced effect of disuse on S, compared to VL, is confirmed by the fact that following immersion, the difference in the transcriptome profile between these muscles became significantly (2 times) smaller than that before it: 1268 and 2493 mRNA, respectively (Fig 1A, Supporting information Table S1).

### Identification of clusters of co-expressed (co-regulated) genes

Using pronounced difference in gene expression between S and VL at baseline and in their response to 6 days of disuse, and an unsupervised approach (the Chinese restaurant process), 39 clusters of co-expressed (co-regulated) genes were identified (Fig. 2A). Then, using hierarchical clustering (Euclidian distance), groups of clusters with similar patterns of gene expression regulation were identified in which *i*) gene expression at baseline differed between muscles (groups comprising 11 and 13 clusters in Fig. 2B) and *ii*) disuse-induced expression changes differed between groups (groups comprising 5, 4, 5, and 9 clusters in Fig. 2C). Then, biological functions of genes in each cluster were examined (functional enrichment analysis) and enriched TFs binding sites (and corresponding TFs) were identified (PWM approach) (Figs. 2A, 3A, and 4A). As a result, using strong cutoff criteria (see Methods), changes in the activity of 239 of 652 TFs expressed in skeletal muscle (TPM>1) were predicted (Supporting information Table S3).

**Fig. 2.**
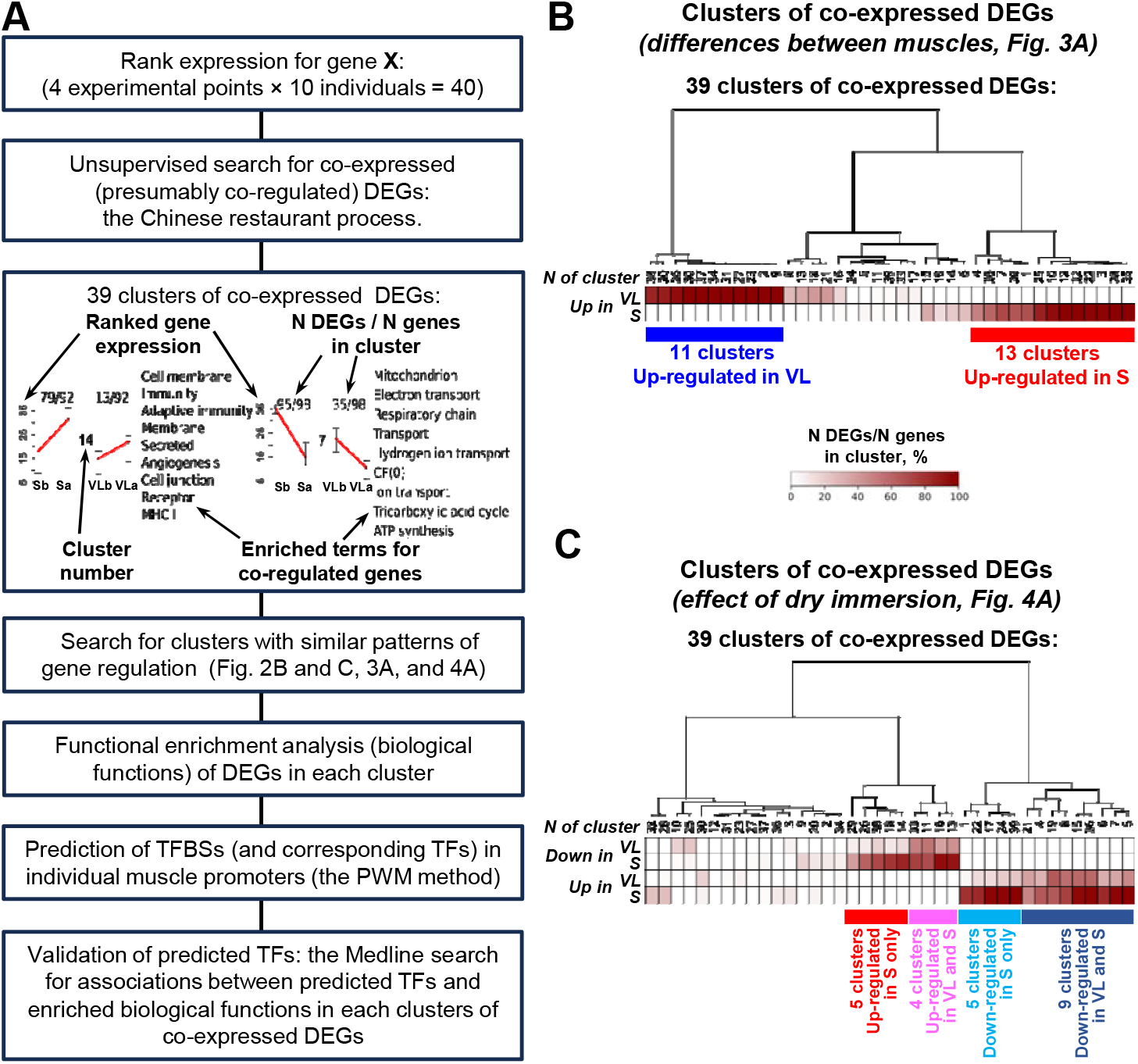
Identification of transcription factors associated with the transcriptome regulation in the “mixed“ *m. vastus lateralis* (VL) and “slow“ *m. soleus* (S) at baseline and after 6 days of disuse. A – The workflow of analysis: identification of clusters of co-expressed (co-regulated) differentially expressed genes (DEGs), transcription factor binding sites (TFBSs) and corresponding transcription factors (TFs) in individual muscle promoters. B and C – Groups of clusters with a similar patterns of gene regulation, in which expression at baseline differs between muscles (B) and disuse-induced expression changes differ between groups (C). The heat map shows the fraction of DEGs in each cluster; groups of clusters and their patterns of gene regulation are indicated by colored rectangles.

**Fig. 3.**
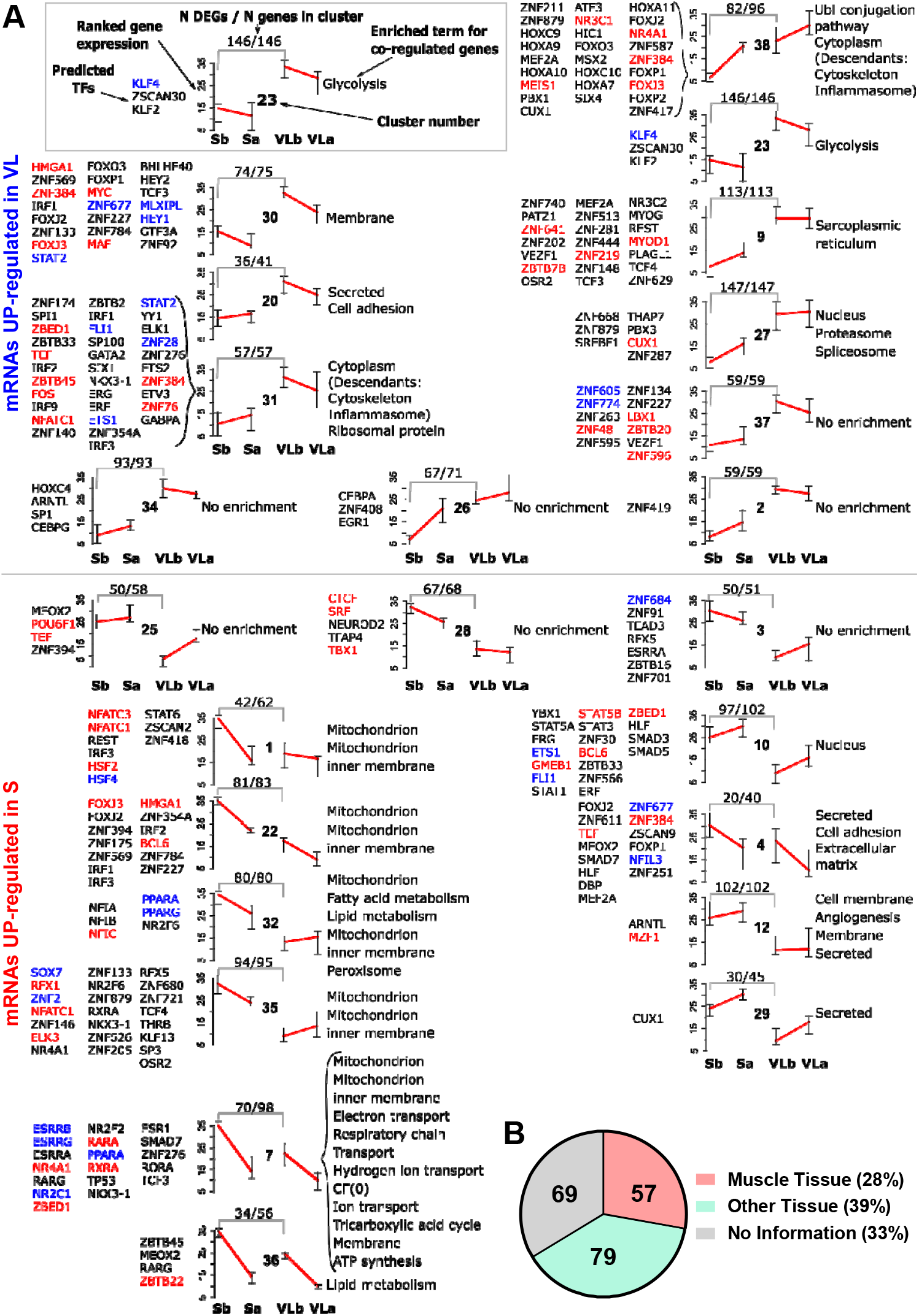
Transcription factors associated with the transcriptome regulation in the “mixed” *m. vastus lateralis* (VL) and “slow” *m. soleus* (S) at baseline (Sb and VLb). A – Predicted transcription factors (TFs), the average level of gene expression (median and interquartile range), and enriched terms are indicated for each cluster of co-expressed (co-regulated) genes (see a legend). Clusters are divided into two groups according to the gene regulation pattern (see Fig 2B). The color of the symbol of each predicted TF corresponds to the pattern of its mRNA expression. B – Validation of predicted TFs: the Medline search for associations between predicted TFs and enriched biological functions in each cluster. The number of validated/not validated TFs is shown.

**Fig. 4.**
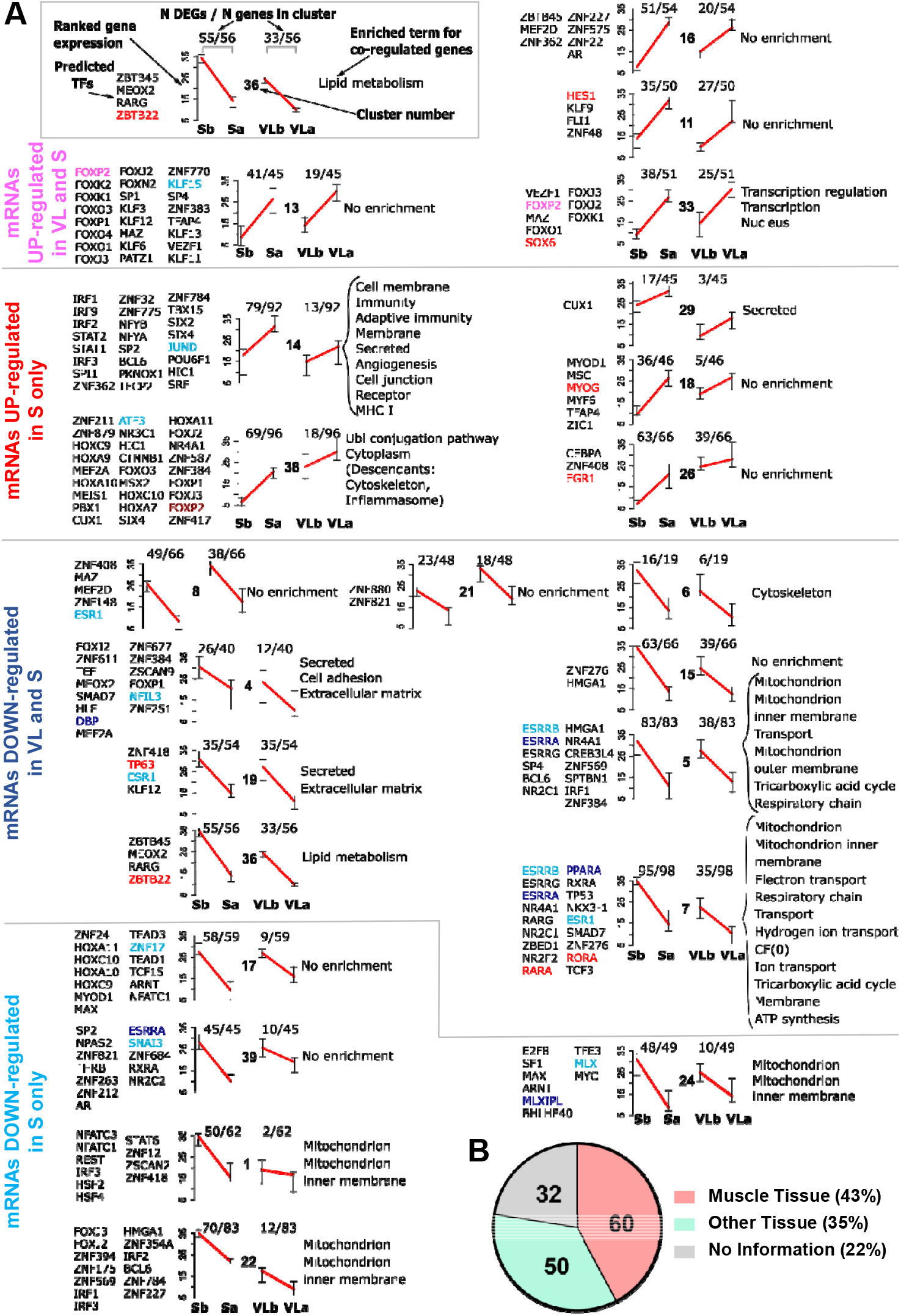
Transcription factors associated with the transcriptome regulation in the “mixed” *m. vastus lateralis* (VL) and “slow” *m. soleus* (S) in response to 6 days of disuse. A – Predicted transcription factors (TFs), the average level of gene expression (median and interquartile range), and enriched terms are indicated for each cluster of co-expressed (co-regulated) genes (see a legend). Clusters are divided into four groups according to the gene regulation pattern (see Fig 2C). The color of the symbol of each predicted TF corresponds to the pattern of its mRNA expression. Sb/a and VLb/a – *m. soleus* and *m. vastus lateralis* before or after 6 days of disuse. B – Validation of predicted TFs: the Medline search for associations between predicted TFs and enriched biological functions in each cluster. The number of validated/not validated TFs is shown.

To assess the efficiency of TF searching in co-expressed genes clusters, an additional analysis was performed to search for TFs associated with genes that changed expression in all experimental conditions (without clustering). In the analysis without clustering, four times fewer TFs were found than in the analysis with clustering: 56 and 239 TFs, respectively; this clearly confirms the advantage of the cluster approach in increasing the depth of regulators prediction.

### Specificity of transcriptome regulation in *m. vastus lateralis* and *m. soleus* at baseline

As expected, in the group of clusters with higher gene expression in S at baseline than in VL, clusters with genes encoding mitochondrial proteins (including oxidative enzymes, mitochondrial ribosomal proteins, etc.; № 1, 7, 22, 32, 35), regulators of lipid metabolism (No. 32 and 36) and angiogenesis (№ 12) were found (Fig. 3A). This is in good agreement with the fact that fatigue-resistant S has an increased mitochondrial and capillary density, as well as the activity and content of oxidative enzymes [12, 13, 16]. Isolation of co-expressed gene clusters allowed to study the specific regulation of various biological functions. Thus, gene clusters with similar functions (e.g. “mitochondria” № 1, 7, 22, 32, and 35 or “fat metabolism” (№ 32 and 36) were associated with different sets of TFs. Among them were both well-studied regulators of mitochondrial biogenesis and lipid metabolism (e.g. TFs of the ESRR, NR4A, PPAR, RAR/RXR families [22, 27]) and a number of TFs whose role in these processes is poorly studied/not studied (Fig. 3A). On the other hand, clusters with higher gene expression in VL than in S included genes encoding glycolytic enzymes (cluster № 23), proteins of the sarcoplasmic reticulum (№ 9), ribosomes (№ 31) and the ubiquitin-proteasome system (№ 27 and 38). This is partly consistent with phenotypic differences: in VL, compared to S, the activity/content of glycolytic enzymes and sarcoplasmic reticulum proteins (calcium metabolism) is increased [12, 13, 16].

### Specificity of disuse-induced transcriptome regulation in various muscles

#### Common regulation for both muscles

Disuse-induced patterns of transcriptome regulation included genes up- or down-regulated in both muscles or in S only (Fig. 2C and 4A). Clusters with genes down-regulated in both muscles were associated with various mitochondrial proteins, regulators of lipid metabolism (clusters № 5, 7, 36), extracellular matrix (№ 4 and 19), and cytoskeletal/sarcomere (№ 6) proteins. Gene expression in these clusters was strongly associated with classical regulators of mitochondrial biogenesis and lipid metabolism (TFs of the ESRR, NR4A, PPAR, RAR/RXR families) [22, 27], as well as with known regulators of extracellular matrix biogenesis and cell adhesion (TFs of the FOX, C/EBP, HOX, SMAD, MEF2, PAR, p53, ESR families from clusters № 4, 19) (see references in Supporting information Table S4). A disuse-induced decrease in the expression of potential target genes indicate a decrease in activity of corresponding TFs. For instance, deactivation of SMAD2/3 was shown to suppress denervation-induced fibrosis in skeletal muscle [28].

Genes up-regulated in both muscles were associated with transcriptional regulation and nuclear proteins (cluster № 33) and TFs belonging to the VEZF1, MAZ, FOX, and SOX families. These factors regulate expression through different mechanisms: VEZF1, by increasing DNA methyltransferase expression, regulates genomic DNA methylation [29], MAZ regulates gene expression by inducing RNA polymerase II arrest and RNA polyadenylation [30], and FOX family TFs can regulate chromatin packing density by changing its accessibility to other TFs [31] (and see references in Supporting information Table S4). SOX TFs, in particular SOX6, are involved in transcription initiation processes, regulation of alternative splicing, and coordinate gene expression via RNA polymerase II stalling [32]. On the other hand, for clusters not associated with biological functions, TFs were also predicted (No. 11, 13, 16) (e.g., belonging to the FOX, KLF, ZNF families), that indicated the presence of common regulation for some genes with different functions.

#### Specific regulation in m. soleus

Genes specifically down-regulated in S were associated with mitochondrial proteins (clusters № 1, 22, 24) (Fig. 4A). Interestingly, these clusters were associated with TFs (NFATC, IRF, HSF, E2F, STAT, MAX/MYC families, etc.) that were different from those predicted for genes down-regulated in both muscles and encoding mitochondrial proteins (see section *Common regulation for both muscles*), that highlighting the specific regulation of gene expression in S. The role of some of these TFs (bHLH-ZIP factors, ARNT, FOXJ3, IRFs, NFATC) in regulating mitochondrial biogenesis has been discussed previously [33] (and see references in Supporting information Table S4). Interestingly, among the predicted TFs were inflammation-induced factors (IRF and STAT families; clusters № 1, 22), which appeared to act as repressors, down-regulating the expression of genes encoding mitochondrial proteins [34, 35] (and see references in Supporting information Table S4).

Importantly, the same TFs were associated with up-regulated genes specific for S (clusters № 14, 38; Fig. 4A) that enriched functional categories related to the immunity, inflammatory response and the ubiquitin-proteasome system, which plays a key role in muscle protein degradation [36]. In addition, up-regulated genes were associated with a number of other TFs (NFY, SP, ZNF, HOX, MEF2A, MEIS1, FOX families, etc.). Some of them (NFY, SP, SIX) was shown to involved in the regulation of inflammatory and immune responses [36] (and see references in Supporting information Table S4). For example, TFs of the NFY family increase the expression of *HLA-DPA1/B1* genes (major histocompatibility complex, class II, DP alpha 1/beta 1) [37], TFs of the HOX family can regulate the chronic inflammatory response through the activation of the key regulator of the inflammatory response – the NF-κB factor [38], transcription factors of the FOX family increase the expression of specific muscle E3 ligases *FBXO32* (*MAFbx*) and *TRIM63* (*MuRF1*), which regulate the ubiquitin-proteasome degradation of sarcomeric proteins [36] (and see references in Supporting information Table S4). Taken together, these results indicate that disuse-induced regulation of gene expression specific for S is closely associated with the inflammatory response and activation of the ubiquitin-proteasome system.

Interestingly, some clusters with genes showing S-specific response to disuse demonstrated difference in expression between S and VL at baseline, indicating that this S-specific response partially explained by the difference in gene expression between the muscles at baseline. Genes in the clusters № 1 and 22 (down-regulated in S only upon disuse) enriched terms related with mitochondrion and mitochondrion inner membrane; genes in the clusters № 26, 29 and 38 (up-regulated in S only upon disuse) – with ubiquitin pathways, inflammasome, and cytoskeleton. These genes (and corresponding TFs) highlight the key feature of S in respondence of disuse.

### Limitations and prospects

Skeletal muscle is composed of various muscle fibers and mononuclear cells, which associated with specific sets of TFs [14]. Myonuclei account for 80-90% of nuclei in skeletal muscle [14, 15], therefore it can be assumed that most of the predicted TFs are related to muscle fibers. Nevertheless, investigating the regulation of specific transcriptomic response to disuse in different skeletal muscle cells using snRNA-seq seems to be a promising direction.

In our work, disuse-induced changes in the expression of genes encoding the predicted TFs (or differences in the expression of these genes between S and VL at baseline) generally did not correlate with the direction of change in TF activity (i.e. with the pattern of regulation of gene expression in clusters of co-expressed genes) (Fig. 3A and 4A). This indirectly confirms that the activity of TFs is weakly regulated at the transcriptional level and emphasizes the prospects of a comprehensive study of post-translational modifications of TFs using mass spectrometry-based analysis.

Promoters of human protein-coding genes contain dozens to hundreds of TF binding sites located adjacent to and overlapping each other; therefore, the expression of each gene is regulated by multiple TFs that bind to DNA transiently (within seconds), compete with or interact with each other, thereby inhibiting or facilitating the binding of other TFs to DNA [39]. Here, a PWM approach with strict cutoff criteria was used, allowing only the most significant (enriched) TF binding sites (and TFs) to be identified. It should be emphasized that in addition to these TFs, many other factors may regulate gene expression at baseline and following disuse. However, experimental validation of such complex regulation of gene expression (involving multiple TFs) is still an unsolved problem even for model cell systems.

Importantly, TF binding sites belonging to the same family are generally very similar. This further highlights the involvement of multiple TFs in the regulation of expression of each gene and means that TFs predicted by the PWM approach correspond to families rather than individual TFs [40].

Despite the above, the validity of TF predictions in this study is indirectly supported by the results of the Medline search (see Methods). According to the literature, two-thirds (one-third for muscle and one-third for non-muscle tissue) of the predicted TFs demonstrated a link with the biological function/s of their corresponding co-regulated genes (Figs. 3B and 4B). The remaining one-third of the predicted TFs appear to be promising candidates for further targeted studies that investigate signaling pathways regulating gene expression in various muscles at baseline and following inactivity.

## Conclusions

As expected, the baseline transcriptomic profiles of the “mixed“ *m. vastus lateralis* and “slow“ *m. soleus* differed significantly. Moreover, these muscles demonstrated a specific transcriptomic response to 6 days of strict disuse. Both muscles demonstrated decreased expression of genes encoding numerous membrane proteins, mitochondria energy metabolism enzymes, and several sarcomeric proteins. However, the specific response of *m. soleus* was significantly more pronounced in terms of the number of gene related to these functional categories, as well as genes associated with the inflammation and the ubiquitin-proteasome system.

Isolation of co-expressed gene clusters allowed to identify different sets of transcription factors associated with the specific regulation of various biological functions in muscles, both at baseline and following disuse. The validity of approximately two-thirds of the predicted transcription factors was indirectly confirmed by analysis of their function described in the literature. These identified transcription factors (especially those that were not previously associated with the regulation of gene expression in skeletal muscle) appear to be promising candidates for future targeted studies that mechanistically investigate gene expression regulation in various muscles at baseline, following disuse or inactivity.

### Data linking

The RNA-seq datasets supporting these findings is available in the NCBI Gene Expression Omnibus (GEO) under accession number GSE271607.

## Acknowledgements

The authors acknowledge the laboratory staffs at IBMP RAS for their excellent assistance in organization of dry immersion experiment.

## Author contributions

OIO, IVR, EST, and DVP conception or design of the work. AAB, PAM, IIP, TFV, IVR, EML, EST, and DVP acquisition, analysis or interpretation of data for the work. AAB, EST, and DVP writing – original draft, review and editing. All authors approved the final version of the manuscript. All persons qualify for authorship are listed.

## Supporting Information

**Supporting Information Table S1**. All and differentially expressed genes (DEGs) between the *m. vastus lateralis* (VL) and *m. soleus* (S) at baseline and after 6 days of dry immersion (DI), as well as DEGs in each muscle after DI.

**Supporting Information Table S2**. Function enrichment analysis for differentially expressed genes between the *m. vastus lateralis* (VL) and *m. soleus* (S) at baseline, as well as for unique and common gene response to dry immersion (DI) in each muscle. The main terms shown in Fig. 1 are depicted in red.

**Supporting Information Table S3**.

***All results of transcription factor binding sites (TFBSs) enrichment analysis***: adjusted fold enrichment (FE)□ and FDR were indicated.

***Significantly enriched TFBSs*** (and 239 corresponding TFs) for groups of clusters with similar patterns of gene expression regulation, where: *i*) gene expression differed at baseline (b) between the *m. vastus lateralis* and *m. soleus* (VL and S, see Fig. 2B) and *ii*) dry immersion (DI)-induced expression changes differed between groups (see Fig. 2C). Significant FE (adjusted fold enrichment >1.5, FDR<0.05) were indicated for individual clusters. Additionally, differentially expressed genes (DEGs between VL and S at baseline (b) and in each muscle after DI) encoding predicted transcription factors are indicated.

A list of ***genes included in each cluster*** is indicated.

**Supporting Information Table S4**.

Validation of predicted transcription factors (TFs): associations of TFs with enriched biological term/function (UNIPROT KW BP/CC) in muscle or other tissues were exanimated in each cluster by a Medline search. For the Medline search, terms: *TF symbol* OR *TF family name* (from the HOCOMOCO v12 database) AND *enriched term for co-regulated genes* AND *muscle* (from Figure 3 and 4) was used. Articles showing a link between predicted TF and enriched biological function in muscle or other tissues were selected.

